# Intra-lineage Plasticity and Functional Reprogramming Maintain Natural Killer Cell Repertoire Diversity

**DOI:** 10.1101/514463

**Authors:** Aline Pfefferle, Benedikt Jacobs, Eivind Heggernes Ask, Susanne Lorenz, Trevor Clancy, Jodie P. Goodridge, Ebba Sohlberg, Karl-Johan Malmberg

**Author notes:** Current address: Department of Haematology and Oncology, University Hospital Erlangen, Erlangen, Germany (Benedikt Jacobs).

## Abstract

Natural killer (NK) cell repertoires are made up of a vast number of phenotypically distinct subsets with different functional properties. The molecular programs involved in maintaining NK cell repertoire diversity under homeostatic conditions remains elusive. Here we show that subset-specific NK cell proliferation kinetics correlate with mTOR activation, and that global repertoire diversity is maintained through a high degree of intra-lineage subset plasticity during IL-15-driven homeostatic proliferation in vitro. High-resolution flow cytometry and single cell RNA sequencing revealed that slowly cycling sorted KIR^+^CD56^dim^ NK cells with an induced CD57 phenotype display increased functional potential associated with inhibitory MHC interactions and activating DAP12 signaling. In contrast, rapidly cycling cells upregulate NKG2A and display a general loss of functionality associated with a transcriptional increase in RNA-binding metabolic enzymes and cytokine signaling pathways. These results shed new light on the role of intra-lineage plasticity during NK cell homeostasis and suggest that the functional fate of the cell is tightly linked to the acquired phenotype and determined by transcriptional reprogramming.

**One Sentence Summary:** High-resolution flow cytometry combined with single-cell RNA sequencing reveal a role for intra-lineage plasticity and functional reprogramming in maintaining phenotypically and functionally diverse NK cell repertoires during IL-15-driven homeostatic proliferation.

## Introduction

Since their discovery in the early 1970s, our view of natural killer (NK) cells has developed from a uniform and short-lived cell population to a group of effectors vastly diverse in terms of phenotype, functionality and life span^1^. Mapping of NK cell subset diversity at the single cell level by mass cytometry revealed that there are more than 10^5^ unique NK cell subsets in the adult human^2^. Much of this diversity results from stochastic expression of killer cell immunoglobulin-like receptors (KIR), but several other heterogeneously expressed activating and inhibitory receptors contribute to the intra and inter-donor diversity of the NK cell repertoire^3^.

The extreme ends of the NK cell differentiation spectrum are well defined, the naïve end consisting of CD56^bright^ NK cells and the mature, terminally differentiated end consisting of CD56^dim^ adaptive, memory-like NK cells^4–6^. CD56^bright^ NK cells are highly responsive to cytokine stimulation and have an immunoregulatory role, while CD56^dim^ NK cells favor receptor ligation input with their function being geared towards cytotoxicity^7^. The CD56^dim^ NK cell subset exhibits large variation in both functionality and phenotype, which can be placed on a spectrum of maturation defined by the expression of key HLA class I binding receptors NKG2A and KIR, as well as CD57, a marker of terminal differentiation^8^. The inhibitory NKG2A receptor is expressed on CD56^bright^ and immature CD56^dim^ NK cells, which can further differentiate by acquiring KIR expression, losing NKG2A expression and acquiring CD57^8^. This, however, is not a strict maturation scheme, as any combination of the above receptors can exist. Nevertheless, phenotyping based on these receptors has served as a useful tool for grouping NK cells based on their functional, metabolic and proliferative ability, as well as their longevity^7,9^. Mouse studies identified critical roles for T-bet and Eomes in the transition from CD27b^+^CD11b^-^ to CD27^-^ CD11b^+^ cells, but the intracellular signaling pathways leading to activation of these transcription factors are still not understood^10,11^. Collin et al., described transcriptional differences between three functional distinct human NK cell subsets (CD56^bright^, CD57^-^CD56^dim^, CD57^+^CD56^dim^) and identified subset specific transcriptional regulators which were shared with the T cell lineage and thus evolutionarily conserved^12^. However, the molecular programs involved in maintaining NK cell repertoire diversity under homeostatic conditions remains elusive.

IL-15 has been identified as the main player in maintaining NK cell homeostasis due to its vital role in survival, development and proliferation^11,13,14^. Immune cells and other non-hematopoietic cells can trans-present IL-15 on the IL-15Rα chain, binding to the IL-15Rβ and the common γ-chain found expressed on the surface of NK cells. The IL-15 complex signals via JAK1/3 activating the transcription factor STAT5. However, the mammalian target of rapamycin (mTOR) can also be activated via IL-15 signaling in a dose-dependent manner^15^. This serine/threonine kinase can form two protein complexes: mTOR complex 1 (mTORC1), which senses nutrients such as glucose and amino acids in the microenvironment, and mTOR complex 2 (mTORC2), which aids in controlling the cytoskeletal organization in the cell^16^. Due to its nutrient sensing ability, mTORC1 controls NK cell metabolism and modifies it accordingly in a process termed metabolic reprogramming. In murine NK cells, mTORC1 activation mediated increased effector function via Granzyme B and IFNγ production by shifting from predominantly oxidative phosphorylation to glycolysis^16–19^. Although mTORC2 has been shown to be sustained by IL-15 induced mTORC1 activation, mTORC2 activation results in a negative feedback loop suppressing mTORC1 induced effector functions by reducing SLC7A5 expression in mice^20^. In humans, the role of mTOR in maintaining homeostasis and as a master regulator for proliferation, education and effector function still needs further investigation.

In humans, the diverse and unique NK cell pool within individuals is well-maintained over time^21^. This stability in receptor repertoires combined with the rapid turnover of NK cells^22^ hints at the important role proliferation plays in replenishing the NK cell pool at steady state. The question arises if this observed stability during homeostatic proliferation is the result of self-renewal from an immature pool of progenitor cells followed by differentiation or arises due to plasticity within the NK cell subsets.

Intra-lineage cell plasticity, also known as functional plasticity, is the term describing phenotypic and functional changes occurring within a given cell lineage^23^. Functional plasticity is an adaptation of the immune system to its surroundings, for examples transition between M1 and M2 phenotypes in macrophages, transition between ILC1, ILC2 and ILC3 or transition between CD4 T helper cells and induced T regulatory cells. Functional plasticity is the result of cytokine or receptor input and is translated into transcriptional changes resulting in modified functionality^24^. NK cell plasticity has largely been unexplored, with the exception of TGFβ-induced NK cell conversion into intermediate ILC1-like cells^25^.

In this study, we show that NK cell repertoire diversity is maintained during IL-15-driven homeostatic proliferation through a combination of mTOR-dependent hierarchy in proliferation capacity and a substantial degree of intra-lineage plasticity. Subset plasticity at the phenotypic level is tightly linked to the functional fate of the cell and associated with upregulation of distinct transcriptional programs that define the acquired phenotype. These results provide new insights into the cellular and molecular programs involved in regulating NK cell homeostasis at the single cell level.

## Results

### Subset specific proliferation kinetics

We set out to study the cellular and molecular events associated with the shift from a quiescent to a proliferative state during homeostatic proliferation in NK cells. To this end we stimulated purified primary human NK cells with 5ng/mL of IL-15 daily, monitored the onset of proliferation and tracked subsequent cell divisions using a proliferation dye. At the population level, cell division began on day 3 and followed a linear path, with roughly one cell division per day (Fig. 1A-B). Slight variation between donors in terms of proliferation and subset distribution was observed (fig. S1). We have previously identified a hierarchy in the proliferative response determined by NK cell differentiation defined through expression patterns of NKG2A, KIR and CD57^8^. Therefore, we stratified the analysis of NK cell proliferation kinetics according to differentiation status into 8 distinct subsets based on the expression of NKG2A, KIR and CD57. CD57 expression had the strongest impact on proliferation speed, resulting in a slightly delayed onset, slower subsequent cell divisions and a low proliferation index (Fig. 1C-D). Notably, NKG2A^-^KIR^+^ NK cells proliferated more slowly than NKG2A^-^KIR^-^ NK cells, independent of CD57 expression (Fig. 1C-D). Importantly, however, irrespective of subset-specific differences in proliferation kinetics, the global NK cell repertoires remained largely stable during the one-week time frame in this experimental set up (Fig. 1E). Thus, this *in vitro* model allowed us to tease apart subset-specific behavior under conditions that mimic natural NK cell repertoire homeostasis.

**Fig. 1.**
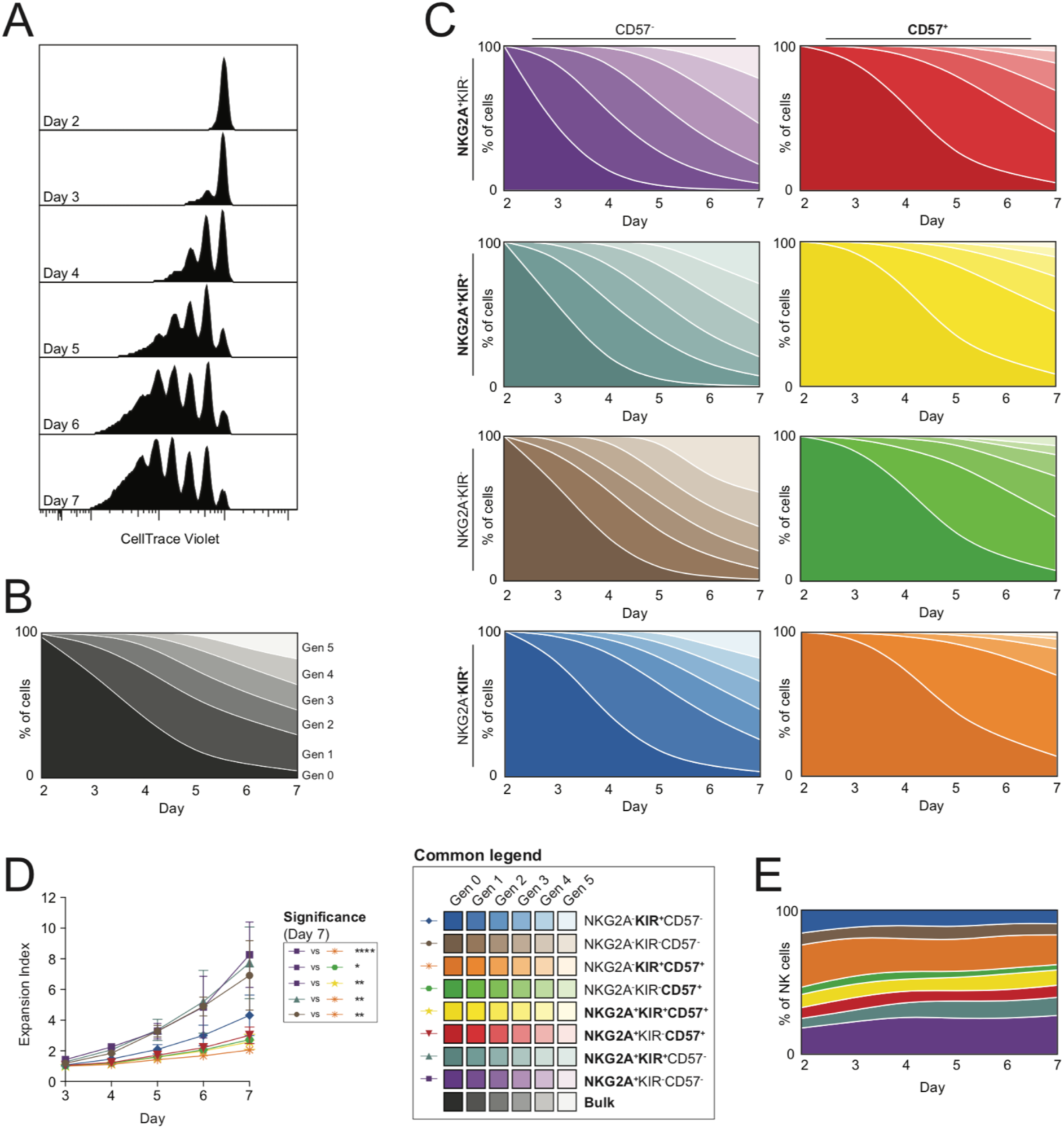
Proliferation kinetics of freshly isolated human NK cell subsets. (**A**) Cell division as measured by CellTrace violet (CTV) dilution for the total CD56^+^ NK cell population from day 2-7 of a representative donor. Each peak represents one generation. (**B** and **C**) Visual representation of the proliferation kinetics of the total CD56^+^ NK cell population (B) and of 8 subsets (defined by Boolean gating of NKG2A, KIR and CD57 expression) (C) in donors from day 2-7. (**D** and **E**) Expansion index (D) and visualization of the same 8 subset frequencies (E) that together comprise the total NK cell population from day 2-7. n = 6-7. Significance was calculated using a Friedman test followed by Dunn’s multiple comparisons test (D). p-values: * <0.05, ** <0.01, *** < 0.001, **** <0.0001.

### mTOR activation determines proliferation kinetics

Given that mTOR has been implicated in NK cell proliferation^15^, we first examined the relationship between mTOR activation and proliferation kinetics. A downstream target of mTORC1 is the ribosomal protein S6 (pS6) which becomes phosphorylated upon mTORC1 activation^15^. We observed an increase in pS6 expression with each additional cell division when comparing different generations in actively cycling cells (day 5) (Fig. 2A-B). Inhibiting mTORC1 in proliferating cells resulted in a decrease in both pS6 and Ki-67 MFI (fig. S2A-B). When looking at the fold change in pS6 expression in bulk NK cells across different days, we observed donor specific upregulation prior to any cell division taking place (day 2) (Fig. 2C). The fold change in pS6 correlated positively with the number of cells having divided on day 3 and followed a similar trend for days 4-5 (Fig. 2C). Early pS6 upregulation was a predictive marker for NK cell proliferation potential in individual donors, as the fold change on day 2 correlated with the percent of cells having divided on day 5 (Fig. 2D).

**Fig. 2.**
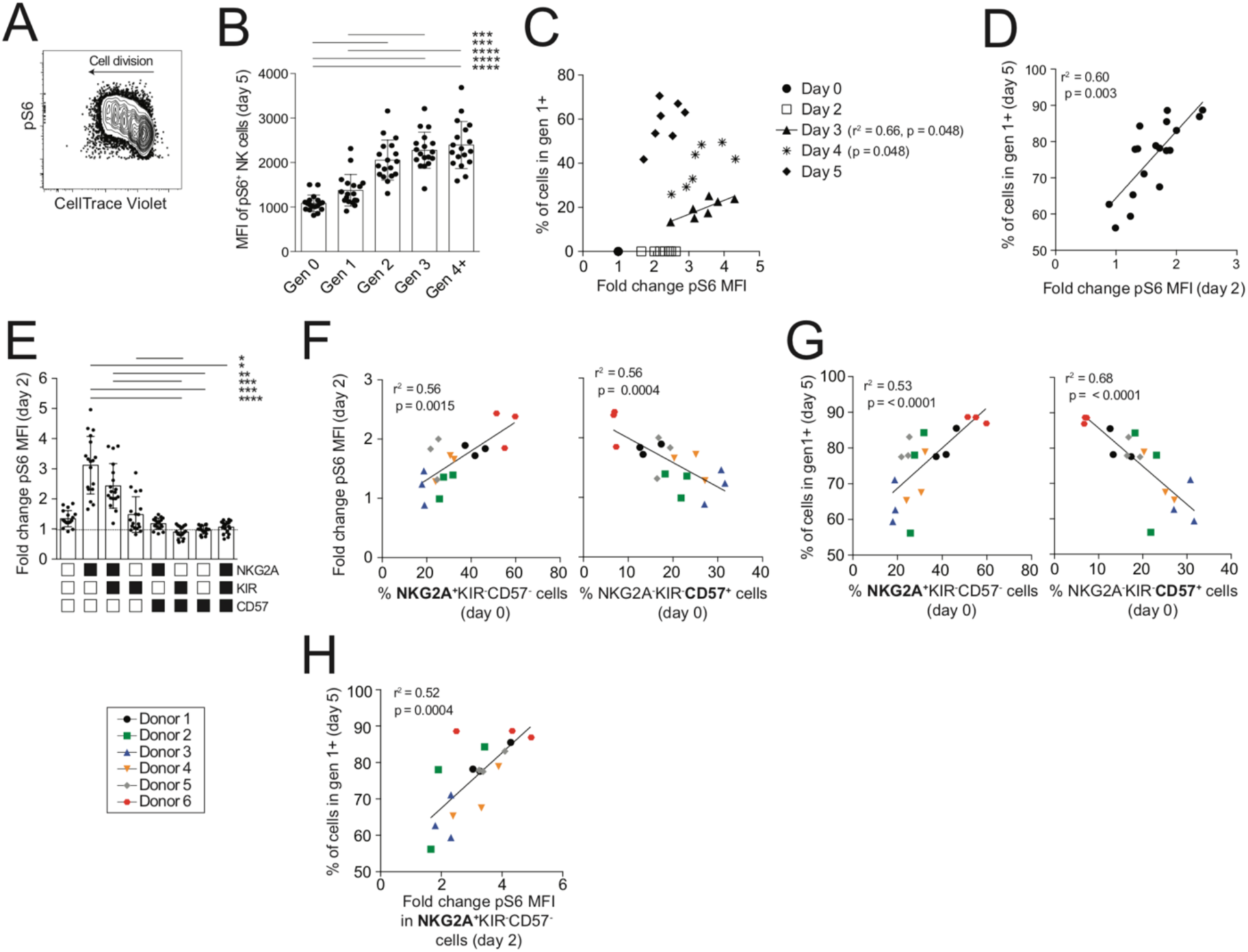
mTOR activation correlates with NK cell proliferation kinetics. (**A** and **B**) Representative FACS plot showing pS6 expression in each generation after 5 days of IL-15 stimulation as visualized by CTV dilution (A) and summary of results (B). (**C**) Fold change in pS6 MFI plotted against the frequency of cells having divided at least once (generation 1+) at baseline (day 0) and on day 2-5. (**D**) Fold change in pS6 MFI on day 2 prior to the onset of proliferation, plotted against the frequency of cells proliferating (generation 1+) on day 5. (**E**) Fold change in pS6 MFI on day 2 within 8 NK cell subsets. (**F** and **G**) Frequency of two subsets (NKG2A^+^KIR^-^CD57^-^ and NKG2A^-^KIR^-^CD57^+^) at baseline plotted against the fold change in pS6 MFI on day 2 for the total NK cell population (F) or the total frequency of cells proliferating (generation 1+) on day 5 (G). (**H**) Fold change in pS6 MFI within NKG2A^+^KIR^-^CD57^-^ cells on day 2 versus the total frequency of cells proliferating (generation 1+) on day 5. In **F**-**H**, each color denotes the same donor sampled monthly over a three-month interval. n = 7-18. Significance was calculated using a Friedman test followed by a Dunn’s multiple comparisons test (B, E) or a Spearman r test (C-D, F-H). p-values: * <0.05, ** <0.01, *** < 0.001, **** <0.0001.

Next, we sampled the same six individuals monthly over a three-month period. We observed a large variation in pS6 fold change in response to IL-15 stimulation among subsets (Fig. 2E). These results prompted us to dissect the role of subset distribution in determining proliferation kinetics at the donor level, focusing on NKG2A^+^KIR^-^CD57^-^ and NKG2A^-^KIR^-^CD57^+^. The percentage of NKG2A^+^KIR^-^CD57^-^ and NKG2A^-^KIR^-^CD57^+^ cells at baseline correlated positively or negatively, respectively, with the fold change in pS6 for the total NK cell population on day 2 (Fig. 2F). Notably, subset distribution at baseline and pS6 fold change on day 2 were both stable within individuals over time, illustrated by the clustering of serial data-points originating from the same individual (Fig. 2F). Furthermore, baseline percentages of NKG2A^+^KIR^-^CD57^-^ and NKG2A^-^ KIR^-^CD57^+^ cells correlated positively or negatively, respectively, with the degree of proliferation observed for the total NK cell population on day 5 (Fig. 2G). Hence, the subset distribution in a given donor correlated with overall pS6 upregulation in that same donor, which in turn correlated with the degree of proliferation at the population level.

A large inter-donor variation in pS6 upregulation was also noted within the NKG2A^+^KIR^-^ CD57^-^ subset, which exhibited the highest fold change in pS6 expression (Fig 2E). Intriguingly, the fold change in pS6 upregulation in the NKG2A^+^KIR^-^CD57^-^ subset prior to proliferation (day 2) correlated positively with subsequent proliferation of all NK cells in that donor (day 5) (Fig. 2H). Therefore, the donor variation in proliferative capacity at a global level is determined in part by the subset composition at baseline and in part by a donor-dependent metabolic set point determining the level of mTOR activation in response to IL-15 stimulation.

### IL-15-induced intra-lineage plasticity

Despite distinct differences in proliferation kinetics and pS6 upregulation between NK cell subsets (Fig. 1D, 2E), the subset frequencies were largely retained at the population level (Fig. 1E), which led us to investigate if transitioning between NK cell phenotypes occurred during IL-15-induced proliferation. To investigate the potential role of intra-lineage plasticity, we sorted three NK cell subsets with distinct differentiation status (NKG2A^+^KIR^-^CD57^-^, NKG2A^-^KIR^+^CD57^-^, NKG2A^-^ KIR^-^CD57^+^) prior to induction of proliferation and monitored the individual cultures for phenotypic diversity after 6 days of IL-15 stimulation (Fig. 3A). Phenotypic stratification of the sorted subsets on day 6 showed a remarkable amount of plasticity, particularly in the NKG2A^-^ KIR^+^CD57^-^ subset, with the most common phenotypic changes being acquisition of NKG2A or CD57 (Fig. 3B). Acquisition of KIR2DL1 or KIR2DL3 expression was less frequent and only occurred in a low percentage of the cells (Fig. 3B).

**Fig. 3.**
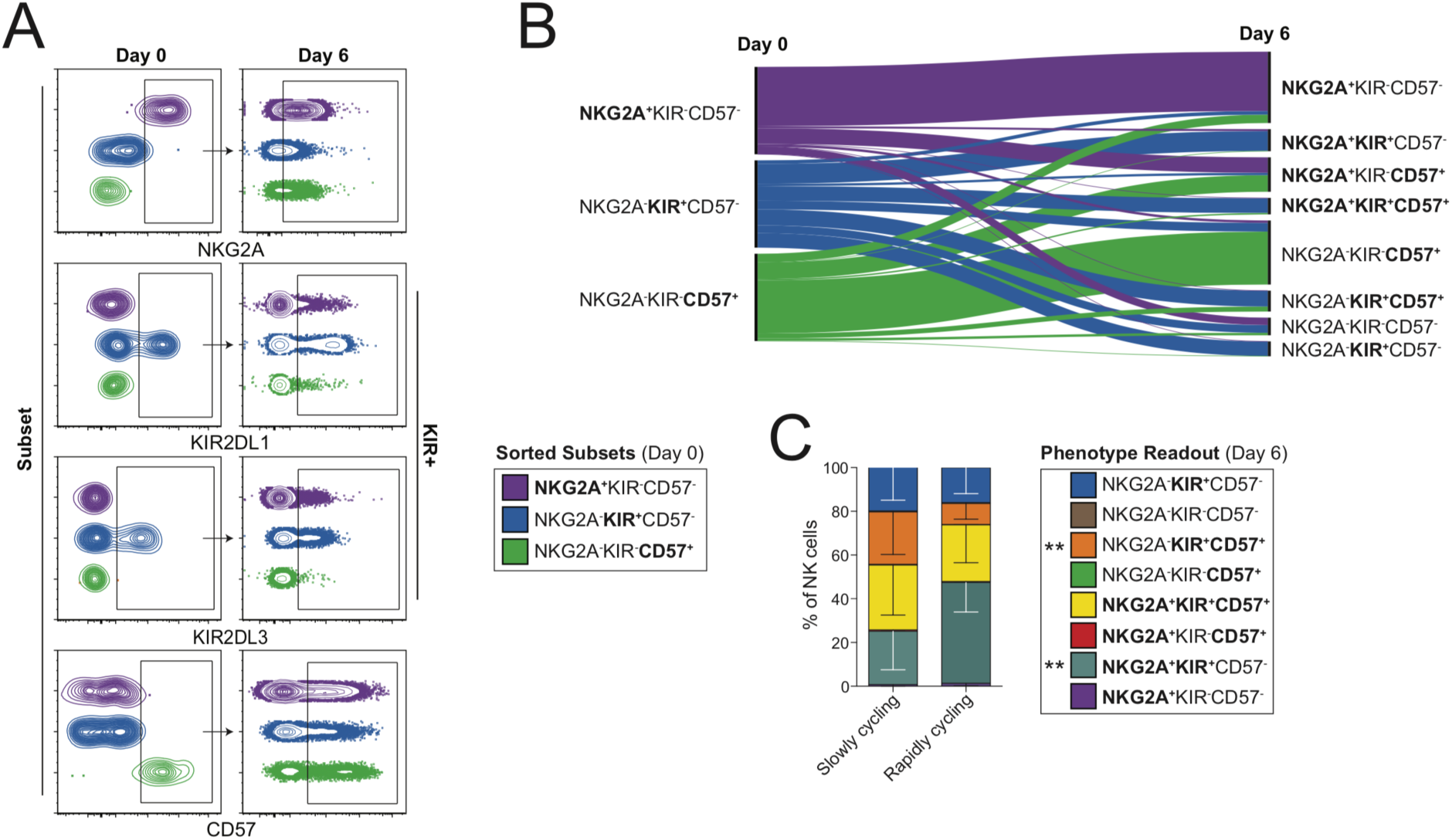
IL-15 induced NK cell intra-lineage plasticity. (**A** and **B**) Representative FACS plot of phenotypic markers (NKG2A, KIR2DL1, KIR2DL3, CD57) used for FACS sorting of three subsets, denoted by colors, on day 0 and the expression of those markers on the sorted subsets after 6 days of IL-15 induced proliferation (A) and visualization of the results (B). (**C**) Distribution of 8 subset frequencies in previously sorted NKG2A^-^KIR^+^CD57^-^ cells further stratified into slowly (Generation 0-1) and rapidly cycling (Generation 2+) cells after 6 days of culture. n=11 donors from 4 independent experiments. Significance of individual subset frequencies between slowly and rapidly cycling cells was calculated using a Wilcoxon signed rank test (C). p-values: ** <0.01. Whiskers show standard deviation.

To determine if the observed plasticity was dependent on the degree of proliferation, we used tSNE analysis to visualize the phenotypic consequences of IL-15 driven plasticity at the generation level (fig. S3). The population map, based on the expression of phenotypic and functional markers, showed a clear distinction between slowly (Generation 0-1) and rapidly cycling (Generation 2+) cells (fig. S3). Applying this stratification based on proliferation kinetics to the NKG2A^-^ KIR^+^CD57^-^ sorted cells, highlighted the impact proliferation kinetics have on the acquired phenotype. High proliferation rate was associated with NKG2A expression while slow proliferation rate was associated with CD57 expression, both within an individual subset (Fig. 3C) and at the population level (fig. S3). Although proliferation impacted the acquired phenotype, the overall degree of plasticity was independent of proliferation, as the size of the original phenotypic subset (NKG2A^-^KIR^+^CD57^-^) was similar in both groups on day 6 (Fig. 3C).

Thus, IL-15-induced plasticity was evident in NK cells across the differentiation spectrum and contributed to the maintenance of repertoire diversity at the population level.

### The acquired phenotype determines functional responses

NK cell differentiation is associated with changes in functional potential^9^. Therefore, we posed the question of whether acquisition of the differentiation markers could modify the cell’s functional capabilities. Acquisition of NKG2A expression in sorted NKG2A^-^KIR^+^CD57^-^ cells resulted in a significantly higher expansion index (Fig. 4A). Conversely, cells that acquired CD57 showed increased ability to degranulate and produce cytokines in response to target cell stimulation (Fig. 4B-C). In line with the functional dichotomy observed in NK cells with induced phenotypes, FACS-sorted slowly cycling cells, exhibiting a higher proportion of CD57^+^ cells, displayed higher killing capacity compared to rapidly cycling, predominantly NKG2A^+^ NK cells (Fig. 4D-E).

**Fig. 4.**
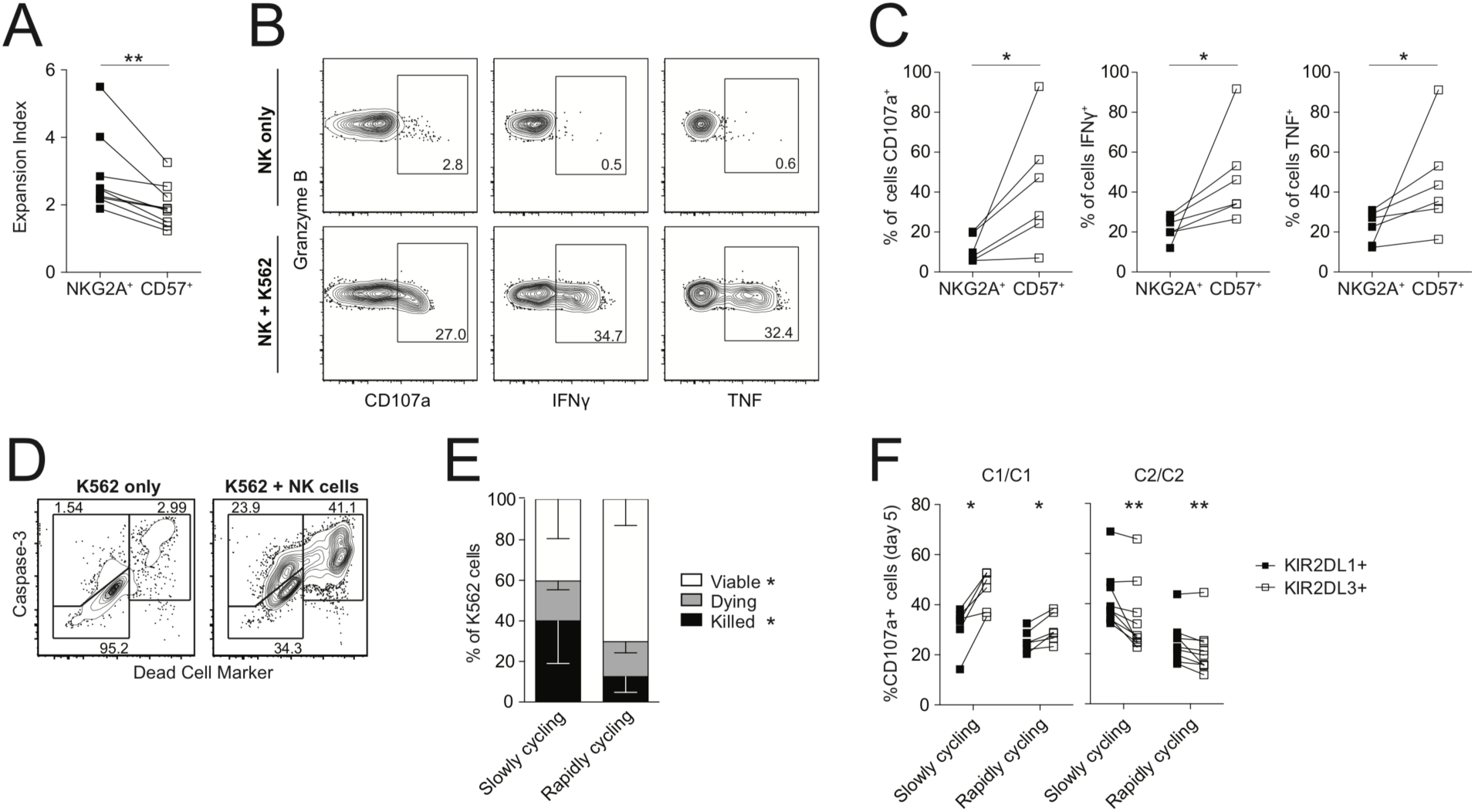
Proliferative and functional consequences of intra-lineage plasticity. (**A)** Expansion index of sorted NKG2A^-^KIR^+^CD57^-^ cells having acquired either NKG2A or CD57 expression on day 6. (**B** and **C**) Representative FACS plots of degranulation (CD107a) and cytokine production (IFNγ^+^, TNF^+^) in controls (NK only) and in response to K562 cells (NK + K562) (B) and summary of results in sorted NKG2A^-^KIR^+^CD57^-^ cells having acquired either NKG2A or CD57 expression on day 6 (C). (**D** and **E**) Representative FACS plots of the K562 killing assay, where K562 cells are stratified into viable (Caspase 3^-^DCM^-^), dying (Caspase 3^+^DCM^-^) and killed (Caspase 3^+^DCM^+^) (D) and summary of results for day 5 NK cells FACS sorted into slowly (Generation 0-1) and rapidly cycling (Generation 2+) cells (E). (**F**) Frequency of CD107a^+^ in single KIR+ NK cells expressing a self or non-self KIR in C1/C1 or C2/C2 donors divided into slowly (Generation 0-1) and rapidly cycling (Generation 2+) cells on day 5. n = 6-9. Significance between NKG2A^+^ and CD57^+^ cells (A and C), slowly and rapidly cycling cells (E) and KIR2DL1^+^ and KIR2DL3^+^ cells (F) was calculated using a Wilcoxon signed rank test. p-values: * <0.05, ** <0.01.

In NK cells, the functional potential is tuned by a processed termed education, where interactions between self-MHC class I ligands and inhibitory receptors are translated into increased effector potential^7^. As functionality was decreased in rapidly cycling NK cells, we wanted to investigate the KIR repertoire during proliferation and examine whether education was inherited in the progeny. t-SNE analysis of the three major inhibitory KIR receptors revealed a narrowing of the KIR repertoire in rapidly cycling cells that mainly expressed one single KIR, while slowly cycling cells were more likely to express multiple KIRs per cell (fig. S4A-B). The education status, determined by the level of degranulation in self and non-self KIR^+^ NK cells in response to target cell stimulation, was inherited in rapidly cycling cells albeit less pronounced compared to slowly cycling cells (Fig. 4F).

These data demonstrate that the loss of functionality observed in rapidly cycling NK cells was mainly related to the induced naïve NKG2A^+^ phenotype compared to the induced mature CD57^+^ phenotype in slowly cycling cells. Hence, the cell’s acquired phenotype, and not its origin, determined its functional capabilities in terms of cytotoxicity and proliferation.

### IL-15 induced transcriptional reprogramming

In order to identify the transcriptional programs associated with the observed phenotypic and functional plasticity we performed single-cell RNA sequencing using sorted NKG2A^-^ selfKIR^+^CD57^-^ NK cells. The cells were cultured with IL-15 for 6 days to allow for plasticity and proliferation to occur, followed by further FACS sorting into slowly and rapidly cycling cells (fig. S5). Additionally, two subsets (NKG2A^+^KIR^-^CD57^-^ and NKG2A^-^selfKIR^+^CD57^+^) were sorted and sequenced at baseline to identify defining transcriptional signatures of these functionally diverse subsets (fig. S5).

To exclude that the differences in proliferation kinetics were due to differential expression of genes involved in cell cycle progression, we clustered all 7556 cells on the expression of 158 cell cycle checkpoint genes and 94 cell cycle genes in our day 6 dataset. Given that the majority of the cells were actively proliferating, t-SNE clustering of NK cells was heavily biased towards cell cycle phase (Fig. 5A). Notably, we found no differences in expression of these cell cycle related genes or checkpoints between slowly and rapidly cycling cells, which were equally distributed in all phases of the cell cycle (fig. S6A-B). As a control, the same set of genes clearly clustered with the stage of the cell cycle (fig. S6A-B). By comparing clusters that differed in cycling speed, but not in cell cycle phase, we could extract differentially expressed genes (DEG) and omit the influence of cell cycle genes. Consequently, we identified 209 DEGs between cluster 2 and 3 (G1 phase) and 617 DEGs between cluster 6 and 8 (G2/M phase), with 130 of these genes being differentially expressed in both phases (Fig. 5B). When the 130 genes were ordered based on the degree of differential expression within each cell cycle phase, we observed a complete overlap in the order of the genes regardless of cell cycle phase analyzed (Fig. 5C).

**Fig. 5.**
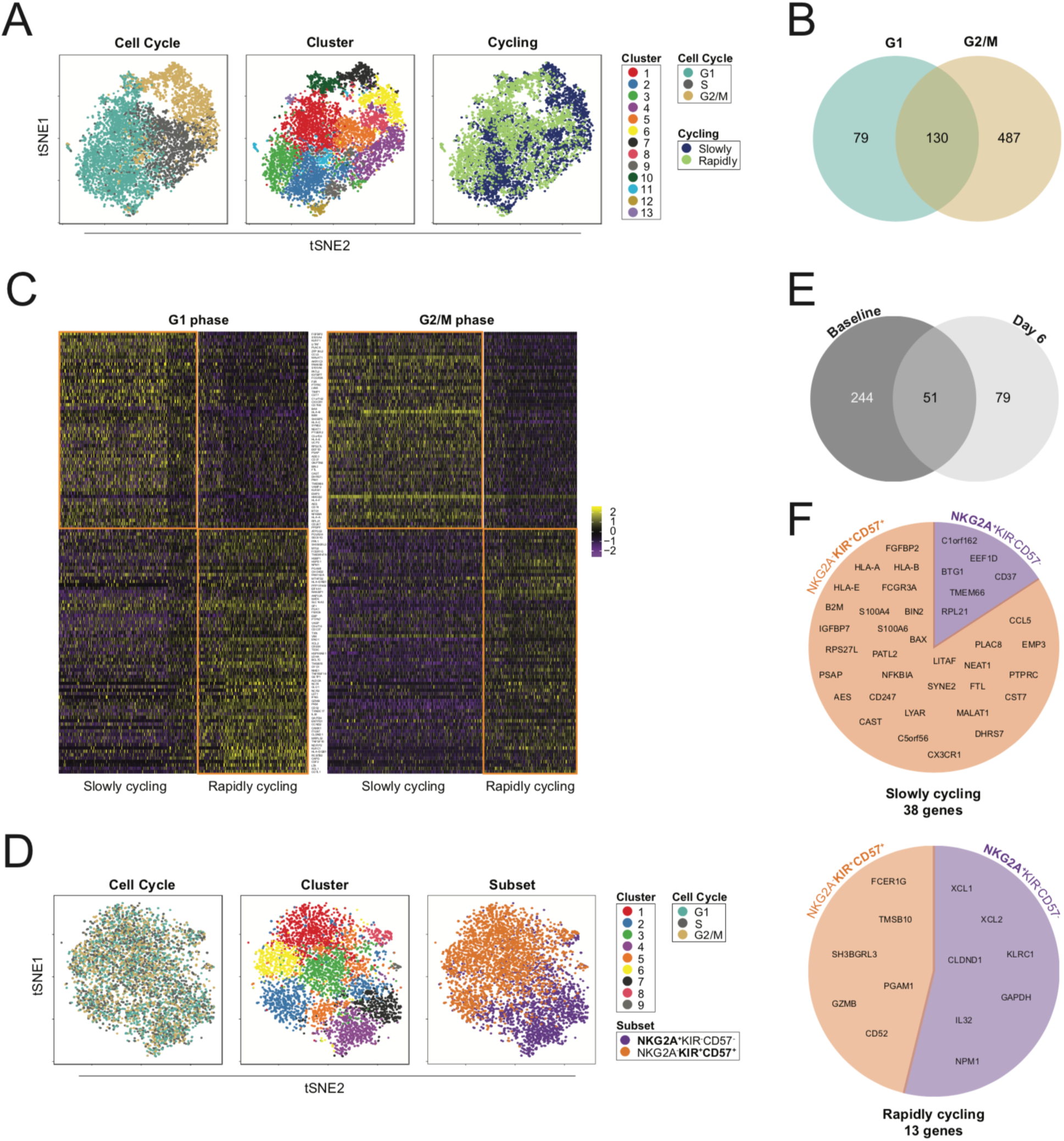
Transcriptional signatures of resting and proliferating NK cell subsets through single-cell RNA sequencing. (**A**) t-SNE plot of single-cell RNA sequencing data of FACS sorted NKG2A^-^selfKIR^+^CD57^-^ NK cells after 6 days of IL-15 stimulation sorted into slowly (Generation 0-1) and rapidly cycling (Generation 2+) cells. (**B**) Venn diagram of differentially expressed genes (DEG) between cluster 2 & 3, representing slowly and rapidly cycling cells in G1 phase, and cluster 6 & 8, representing slowly and rapidly cycling cells in G2/M phase. (**C**) Expression heatmap of the 130 commonly DEGs in both G1 and G2/M phases of the cell cycle. (**D**) t-SNE plot of single-cell RNA sequencing data of FACS sorted NKG2A^+^KIR^-^CD57^-^ and NKG2A^-^ selfKIR^+^CD57^+^ resting NK cells at baseline. (**E**) Venn diagrams of the 295 DEGs between NKG2A^+^KIR^-^CD57^-^ (cluster 4+7) and NKG2A^-^selfKIR^+^CD57^+^ (cluster 1+6) cells at baseline and the 130 DEGs between slowly and rapidly cycling cells at day 6. (**F**) Pie chart of DEGs commonly upregulated in either slowly or rapidly cycling cells, stratified based on the subset it was upregulated in at baseline (NKG2A^+^KIR^-^CD57^-^ or NKG2A^-^KIR^+^CD57^+^). n = 1-2.

To address whether the IL-15 induced subset plasticity was associated with transcriptional reprogramming we first identified genes that were differentially expressed between mature NKG2A^-^selfKIR^+^CD57^+^ and naïve NKG2A^+^KIR^-^CD57^-^ NK cells at baseline. Single-cell RNA sequencing revealed 295 DEGs between clusters most clearly representing the NKG2A^+^KIR^-^ CD57^-^ and the NKG2A^-^selfKIR^+^CD57^+^ subset (Fig. 5D and fig. S6C). Next, we compared the 130 DEG of the induced subsets (day 6) with the 295 DEG between the corresponding subsets at baseline and identified 51 overlapping genes (Fig. 5E). Of these 51 differentially expressed genes, 38 were upregulated in slowly cycling cells and only 13 in rapidly cycling cells (Fig. 5F). 84% of the slowly cycling gene signature was shared with the mature NKG2A^-^selfKIR^+^CD57^+^ signature at baseline, while 54% of the rapidly cycling gene signature was shared with the naïve NKG2A^+^KIR^-^CD57^-^ signature at baseline (Fig. 5F). Hence, IL-15-driven intra-lineage plasticity and the associated shift in functional responses was associated with transcriptional reprogramming towards the baseline signature of NK cells with the acquired phenotype.

### Cytokine-versus receptor-driven biological programs

Although the functional potential of NK cells at different differentiation stages is well documented, the transcriptional signatures mediating these functionalities are not as well defined. Pathway analysis of the 130 DEGs showed a clear difference in potential upstream regulators between slowly and rapidly cycling cells (Table 1). Upstream regulators of rapidly cycling cells included cytokines (IL-15, IL-18), transcription regulators (MYC, HIF1A, STAT5A, JUN), stimulatory agents (PMA, ionomycin), as well as activation induced kinases (AKT1) and complexes (NFkB). Conversely, upstream regulators of slowly cycling cells included an array of inhibitory molecules (cyclosporin A, sirolimus, wortmannin, prostaglandin E2, PD98059, trichostatin A) and cytokines (IL-6, IL-27). Intriguingly sirolimus, better known as rapamycin, is a potent inhibitor of mTORC1, which we have identified as the main determinant of IL-15 induced proliferation. Furthermore, a number of the upstream regulators of rapidly cycling cells are associated with mTOR signaling, including mTOR phosphorylation via AKT1^27^.

**Table 1.** Top upstream regulators of DEG between slowly and rapidly proliferating NK cells. Activation z-score, p-value and molecular type of selected upstream regulators determined using Ingenuity Pathway Analysis of differentially expressed genes between slowly (Generation 0-1) and rapidly cycling (Generation 2+) cells.

|  | Upstream Regulator | Molecule Type | Activation z-score | p-value of overlap |
| --- | --- | --- | --- | --- |
| Slowly cycling | NLRC5 | transcription regulator | 2,758 | 5,43E-11 |
|  | cyclosporin A | calcineurin inhibitor | 2,657 | 9,42E-15 |
|  | IFNA2 | cytokine | 2,468 | 5,19E-17 |
|  | IL6 | cytokine | 2,229 | 1,48E-12 |
|  | sirolimus | mTOR inhibitor | 2,071 | 2,63E-12 |
|  | wortmannin | PI3K inhibitor | 1,757 | 1,45E-10 |
|  | prostaglandin E2 | chemical - endogenous mammalian | 1,738 | 2,97E-13 |
|  | PD98059 | MAPK inhibitor | 1,669 | 2,62E-15 |
|  | trichostatin A | HDAC inhibitor | 1,659 | 4,02E-11 |
|  | IL27 | cytokine | 1,353 | 2,75E-16 |
| Rapidly cycling | MYC | transcription regulator | -2,991 | 7,23E-13 |
|  | ionomycin | chemical reagent | -2,920 | 2,85E-11 |
|  | HIF1A | transcription regulator | -2,832 | 6,12E-13 |
|  | phorbol myristate acetate | chemical drug | -2,471 | 1,68E-17 |
|  | IL18 | cytokine | -2,063 | 1,08E-13 |
|  | AKT1 | kinase | -1,950 | 1,71E-12 |
|  | STAT5A | transcription regulator | -1,937 | 4,04E-19 |
|  | NFkB (complex) | complex | -1,896 | 1,18E-13 |
|  | JUN | transcription regulator | -1,868 | 3,05E-11 |
|  | IL15 | cytokine | -1,630 | 1,22E-38 |

We applied Gene Ontology’s gene over-representation analysis, looking at molecular function, biological processes and pathways significantly different between slowly and rapidly cycling cells (Fig. 6). The common theme for slowly cycling cells was receptor-based interactions leading to increased cytotoxicity, as shown by antigen presentation, DAP12 signaling, IFN signaling, calcium homeostasis and the endosome/vacuolar pathway. Genes that highly contributed to these pathways were the MHC class I molecules (HLA-A/B/C/E/F), while B2M and CD16 (FCGR3A) also moderately contributed. In contrast, rapidly cycling cells were associated with increased cytokine activity and signaling as well as increased metabolic activity (glycolysis, gluconeogenesis).

**Fig. 6.**
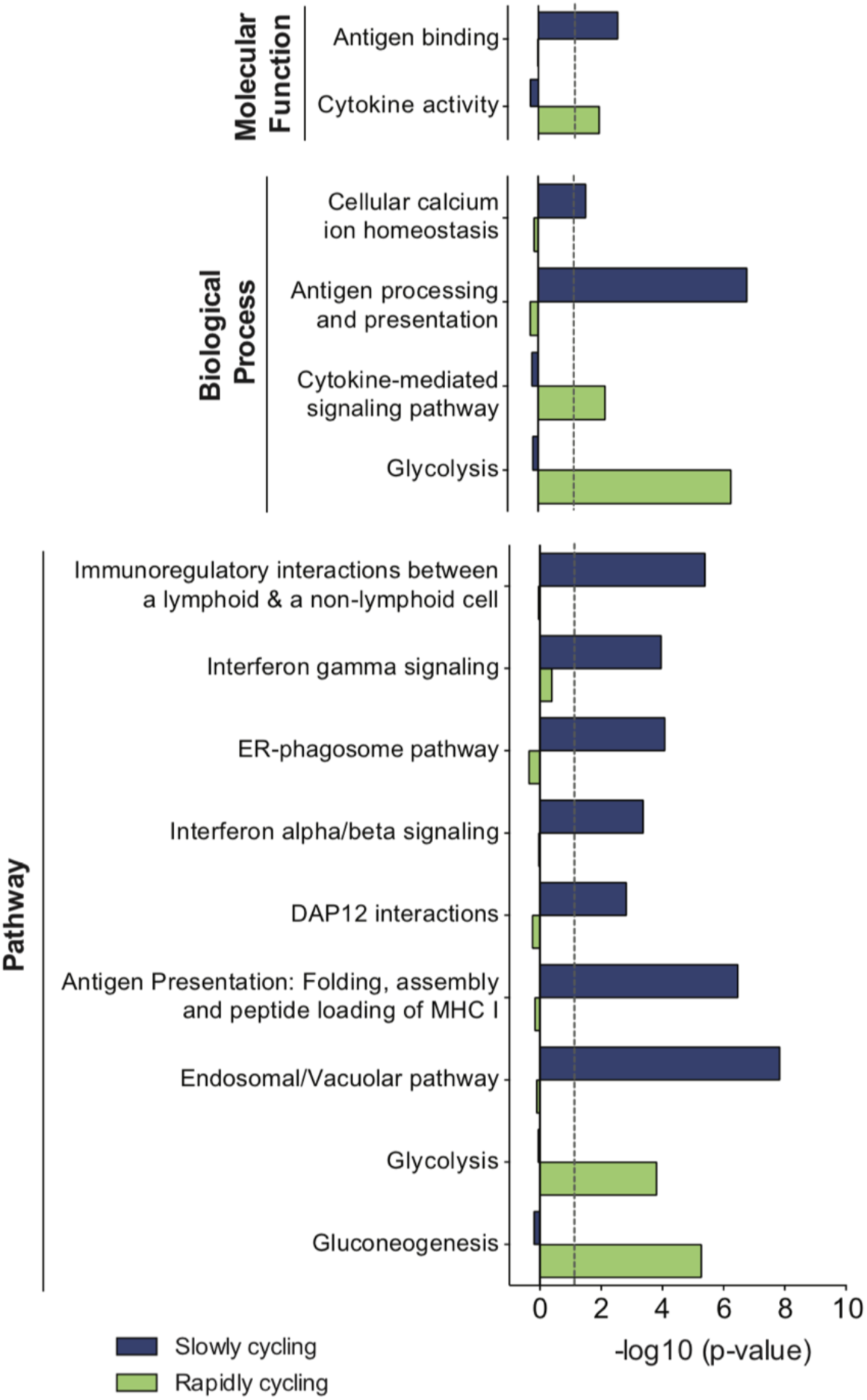
Gene over-representation analysis of 130 differentially expressed genes between slowly and rapidly cycling cells. Selected GO terms for molecular function biological process and pathways that were significantly differentially represented in slowly (Generation 0-1) versus rapidly cycling (Generation 2+) cells. n = 2. Significance was calculated using Fisher’s Exact test followed by false discovery rate correction of the p-value. A negative log_10_ value of the false discovery rate adjusted p-value > 1.3 is deemed significant.

Our findings revealed stark differences in the molecular signature of slowly and rapidly cycling cells, hinting at increased metabolism and activation leading to cytokine dependence in rapidly cycling cells. Slowly cycling cells, on the contrary, acquired transcriptional signatures of mature NK cells with a focus on receptor-based interactions, potentially explaining the increased functionality upon target cell interactions.

## Discussion

IL-15 is a key homeostatic cytokine for NK cells and has been linked to vital processes including survival, differentiation, functionality and proliferation^14,15,17,28–30^. However, it remains elusive how these processes come together at the population level to maintain stable and yet highly diversified subset repertoires within individuals. We developed an *in vitro* model to study functional consequences and transcriptional programs involved in IL-15 driven homeostasis and subset plasticity at the single cell and repertoire level. We noted a striking mTOR-dependent hierarchy in proliferative responses among discrete NK cell subsets. Yet, despite profound subset-specific differences in proliferation kinetics, the global repertoires remained relatively stable suggesting that proliferation differences between subsets could be balanced by plasticity.

Intra-lineage plasticity, defined as phenotypic and functional changes occurring within a given cell lineage,^23^ remains largely understudied in the context of NK cells. Most studies published so far have examined dynamic phenotypic changes in the NK cell repertoire in a tumor setting, either using mouse models or stimulation with tumor cell lines^25,31^. Herein, we observed a high degree of NK cell plasticity across the differentiation spectrum in response to IL-15 stimulation. We focused our analysis on NKG2A^-^KIR^+^CD57^-^ NK cells, a subset at an intermediate stage of differentiation, and followed their fate after FACS sorting and subsequent IL-15-induced proliferation. At the phenotypic level, we noted a high degree of plasticity with the dominant induced phenotypes being cells that retained KIR expression and acquired either NKG2A or CD57 expression. These two major phenotypes were linked to different functional fates, with induced NKG2A^+^KIR^+^CD57^-^ NK cells displaying strong proliferative responses, whereas induced NKG2A^-^KIR^+^CD57^+^ NK cells exhibited increased responsiveness to target cell stimulation.

To date, studies looking at homeostatic proliferation have largely been carried out in lymphopenic mice with a focus on T cell proliferation^54–56^. Spontaneous proliferation occurred in severely lymphopenic mice, hallmarked by rapid cell division (<24h) without the need for cytokine stimulation, resulting in an acquired memory phenotype, diversification of the T cell receptor repertoire and cytokine production upon stimulation. Homeostatic proliferation, on the other hand, was observed in mildly lymphopenic hosts and at a slower division rate, with proliferation requiring both TCR interaction and cytokine stimulation (IL-7). Target cell lysis by CD8^+^ T cells was only observed by day 12 post transfer, highlighting a functional inability during the initial phase of proliferation. Most importantly, once the proliferation cues were removed, the proliferating cells reverted back to their initial phenotype whereas this reversal was not observed in RAG^-/-^ hosts due to continued proliferation^55^. These studies bear striking similarity with the functional dichotomy between rapid and slowly cycling NK cells and it would be interesting to determine if rapidly cycling cells could regain functionality in the absence of proliferation cues.

To address whether the phenotypic and functional shift observed in our *in vitro* model was associated with transcriptional reprogramming, we projected the differentially expressed genes in cells with the induced phenotypes onto baseline transcriptional signatures of naïve and differentiated NK cells. Since we did not sort NKG2A^+^KIR^+^CD57^-^ at baseline, the induced NKG2A^+^KIR^+^CD57^-^ phenotype in rapidly cycling cells had to be related to transcriptional signature of the more naïve NKG2A^+^KIR^-^CD57^-^ cells. Slowly cycling cells had a transcriptional signature that overlapped to a great degree with more differentiated NKG2A^-^selfKIR^+^CD57^+^ cells at baseline. Although rapidly cycling cells showed increased expression of genes defining naïve NKG2A^+^KIR^-^CD57^-^ cells, they also exhibited increased expression of genes associated with more mature cells, such as GZMB, likely reflecting an IL-15 primed state. Altogether, these results suggested that IL-15 induced intra-lineage plasticity was associated with transcriptional and functional reprogramming, resulting in two functionally diverse NK cell populations.

One of the key regulatory mechanisms behind immune cell differentiation and function is metabolism, at the center of which lies mTOR^32^. mTOR controls an important checkpoint for both activation and differentiation in NK cells, but the exact signaling cascade upstream and downstream of mTOR mediating this effect is unknown^16,20,30,33–35^. Studies in mice have identified a threshold of IL-15 stimulation which needs to be surpassed for mTORC1 activation to occur, while suboptimal stimulation resulted in STAT5 activation only. Similarly, we observed a large variation in mTORC1 activity, both at the donor level but more intriguingly at the subset level where the level of activation correlated with and could predict the subsequent degree of proliferation. Naïve NKG2A expressing cells are highly responsive to cytokine stimulation^36^. In line with this notion, rapidly cycling NKG2A^+^ NK cells exhibited the highest fold change in pS6 expression, suggesting that they reached the threshold needed for mTORC1 activation. Hence, mTOR activation may play a role in inducing the naïve NKG2A^+^KIR^+^CD57^-^ phenotype in cytokine responsive, rapidly cycling cells. In contrast, sirolimus, an mTOR inhibitor^37^, was one of the top upstream regulators of the gene program associated with the induced mature NKG2A^-^ KIR^+^CD57^+^ phenotype. These data suggest that the mTOR pathway is most important during early differentiation whereas terminal differentiation may be regulated by other IL-15 induced pathways.

Single-cell RNA sequencing revealed a prominent IL-15-driven metabolic reprogramming in rapidly cycling cells, with significantly increased expression of glycolytic enzymes including PGK1, ENO1, PGAM1, ALDOA, GAPDH, GPI, LDHA and PKM. These signatures match the IL-15 induced metabolic reprogramming observed in mouse studies which are dependent on mTOR activation^15–17^. Metabolic reprogramming is necessary to meet the energetic and biosynthetic needs of proliferating cells. Stimulation induced aerobic glycolysis allows for increased glycolytic flux combined with increased rates of biosynthesis^38^. Recently a new functional role for selected metabolic enzymes has been described, namely regulating transcription and translation by translocating to the nucleus and binding to RNA and DNA^39–41^. Included in this group are GAPDH, ENO1, PKM, LDHA and GPI. In T cells, during high glycolytic flux, GAPDH has been shown to induce translation of IFNγ and IL-2 while it’s interaction with Rheb is inhibited, releasing Rheb to activate mTORC1^40,42^. Similarly, PKM2, one of two transcripts of PKM, can activate mTORC1 by phosphorylating another inhibitor AKT1S1, which could account for increased mTORC1 activation in rapidly cycling cells^43^. GPI, yet another glycolytic enzyme, has been shown to promote growth, increase motility of fibroblasts in a tumor setting, and can be inhibited by the insulin growth factor binding protein 3 (IGFBP3)^40^. Although IGFBP3 was not differentially expressed in our dataset, slowly cycling cells did have significantly increased IGFBP7 expression as well as decreased GPI expression. Furthermore, de Rosa et al have identified an important role for enolase-1 (ENO1) in transforming highly metabolically active and proliferating conventional T cells into iTreg cells^44^. This complete reversal of function of the T cell was mediated by ENO1 translocating to the nucleus. Evidently, activation induced metabolic reprogramming in rapidly cycling cells could account for the functional differences we observe in this population.

Rapidly cycling cells also exhibited increased expression of genes associated with actin filament organization^45^. These genes included COTL1, CAPG, ALDOA, TMSB10, VIM, and VASP, where COTL1 and CAPG are two of the most highly differentially expressed genes (data not shown). Considering the role actin plays in immune synapse formation, conjugate formation and the transportation of cytotoxic granules to the surface^46^, modified actin polymerization during intense proliferation could result in a number of effects that impact target cell responsiveness. Although IL-15 stimulation induced granzyme B production in a proliferation dependent manner, this did not translate into increased target cell killing. COTL1 and CAPG are genes normally only expressed in bright NK cells (data not shown), which similar to the rapidly cycling cells studied here, are highly responsive to cytokine input, have a high proliferative capacity and reduced cytotoxic potential. Hence, further studies looking at the role of actin organization and immune synapse formation in rapidly cycling cells is warranted. It will be important to delineate the mechanisms that allow the cell to switch from a proliferative mode to a cytotoxic and target-seeking mode.

The gene signature contributing to the pathway analysis in slowly cycling cells was dominated by MHC class I molecules (HLA-A/B/C/E/F), B2M and FCGR3A expression. This is in line with the flow cytometry data, as the more differentiated slowly cycling cells commonly expressed multiple KIRs per cell. Although HLA-KIR interactions normally occur in *trans, cis* interactions have been observed in mice^47,48^. Ly49 receptors have a flexible stalk, which KIR do not, allowing for a *cis* interaction to take place^49^. Nonetheless, HLA-LILR *cis* interactions have been shown in humans and therefore *cis* HLA-KIR interactions cannot be excluded^50,51^. In mice, higher expression of MHC on the NK cells’ surface led to increased MHC-Ly49 binding, effectively restricting the number of Ly49 molecules available to interact with potential target cells, and thus lowering the activation threshold required^47,52,53^. We found that the increased inhibitory interactions combined with FCGR3A expression was associated with increased cytotoxicity in slowly cycling cells. Similarly, a lack of inhibitory signaling observed in rapidly cycling cells due to limited HLA-E expression could turn the cells hyporesponsive. Intriguingly, while MHC class I molecules were increased in slowly cycling cells, two MHC class II molecules were increased in rapidly cycling cells, namely HLA-DRB1^31^ and HLA-DQB1, potentially indicating an activation signature due to cytokine responsiveness.

Our results provide new insights into how, during homeostasis, intra-lineage plasticity in response to IL-15 effectively maintains phenotypically and functionally diverse NK cell repertoires within individuals. Exploring the role plasticity plays in tissue-specific niches or in conditions of perturbed homeostasis, such as during infection or post stem-cell transplantation, will further our understanding of the mechanisms involved in the diversification and differentiation of NK cells.

## Materials and Methods

### Cell processing and proliferation protocol

Using density gradient centrifugation (Ficoll-Hypaque; GE Healthcare), peripheral mononuclear cells (PBMC) were isolated from healthy blood donors obtained from the Karolinska University Hospital or Oslo University Hospital Blood bank (dnr: 2016/1415-32). NK cells were further isolated using magnetic-activated cell sorting (MACS) (NK cell isolation kit; Miltenyi Biotech) and labeled with 1μM cell proliferation dye CellTrace Violet (CTV; Life Technologies) or Carboxyfluorescein succinimidyl ester (CFSE; Life Technologies). The labeled NK cells were resuspended in Stem Cell Growth Medium (SCGM; CellGenix) supplemented with 10% human serum (Sigma), 2mM L-glutamine (GE Healthcare) and 5ng/mL IL-15 (R&D) at 1,5×10^6^ cells/mL and cultured in 96 U-bottom wells in a final volume of 200μL at 37°C and 5% CO_2_. IL-15 was replenished daily and 100μL of medium was replaced with fresh medium on day 3 of culture. Visualization plots for proliferation (Fig. 1B-C, E) and plasticity (Fig. 3B) were created using RAW Graphs^57^.

### Flow cytometry

Cells were harvested and stained in 96 V-bottom plates using the following fluorochrome conjugated antibodies (clone names in brackets): CD107a-AF700 or PE-Cy5 (H4A3), CD14-V500 (MφP9), CD19-V500 (HIB19), CD3-V500 (UCHT1), CD57-BV605 (NK-1), Granzyme B-AF700 (GB11) from Beckton Dickinson; CD158a,h-PE-Cy7 (EB6B), CD158b1/b2,j-PE-Cy5.5 (GL183), CD159a-APC-AF750 (Z199), CD3-PE-Cy5 (UCHT1), CD56-ECD (N901) from Beckman Coulter; CD158e1-APC or Biotin (DX9), CD16-BV785 (3G8), CD57-FITC (HNK-1), IFNγ-BV785 (4S.B3) from Biolegend; CD158a-APC or APC-Vio770 (REA284), CD158e/k-Biotin or PE (5.133), CD159a-PE-Vio770 (REA110), CD159c-PE (REA205), Ki-67-PE Vio770 (REA183), KIR2D-Biotin or PE (NKVFS1) from Miltenyi Biotec; TNFα-APC (MAb11) from eBioscience; p-S6 ribosomal protein (S235/236)-PE (D57.2.2E) from Cell Signaling Technologies; KIR2DL3-FITC (180701) from R&D Systems. Viability was determined using LIVE/DEAD Fixable Aqua Dead Cell Stain kit (DCM) for 405 nM excitation and biotin-conjugated antibodies were visualized using Streptavidin-QD705 (Life Technologies). Cells were fixed and permeabilized using Fixation/Permeabilization Kit (eBioscience) followed by intracellular staining.

Samples were acquired on an LSR-Fortessa equipped with a 355, 405, 488, 561, and a 639 nm laser. Sorting experiments were performed either on a FACSAria Fusion (Becton Dickinson) equipped with a 405, 488, 561 and a 633 laser or on a FACSAriaII (Becton Dickinson) equipped with a 405, 488, 561 and 640 nm laser. Cell sorting was performed at 4°C. Data was analyzed in FlowJo version 9 (TreeStar, Inc.).

To minimize variation for daily readouts over the course of one week, CS&T beads (BD Bioscience) and application settings in FACS DIVA were used to eliminate daily fluctuations in the LSR-Fortessa. Additional controls included usage of a reference sample, consisting of a buffy coat frozen in small aliquots which was thawed and stained for each experimental readout. Values were then adjusted based on the reference values which were normalized over the days.

Fold change of pS6 expression on a particular day was calculated by comparing the observed expression to baseline expression at day 0. Expansion index, the overall fold expansion of the culture, was calculated based on the CTV dilution using the Proliferation tool in FlowJo.

t-distributed Stochastic Neighbor Embedding (t-SNE) analysis was performed using the Rtsne package (http://cran.r-project.org/package=Rtsne) in R version 3.1.0. Events from one representative donor were divided into generations by manual gating on CTV in FlowJo version 9. Then, 20 000 events were randomly sampled from each generation and data was pooled and arcsinh transformed using cofactor 150. The t-SNE calculations were based on the markers NKG2A, CD57, CD16, KIR2DL1, KIR2DL1/S1, KIR2DL3, KIR2DL2/S2/L3, KIR3DL1, Granzyme B and pS6. Plots were generated using FlowJo or the ggplot2 R package (http://ggplot2.org) and red borders were added using Photoshop CS6 (Adobe). “nKIR” values were generated based on manual gating of KIR2DL1, KIR2DL3 and KIR3DL1 in FlowJo.

### Functional assays

Cells were harvested, counted and seeded at an effector:target ratio of 1:1 in complete RPMI medium (10% Fetal calf serum, 2mM L-glutamine). For the natural cytotoxicity and the antibody-dependent cellular cytotoxicity (ADCC) assay the target cells were K562 and P815 pre-incubated with purified CD16 antibody (3G8, Becton Dickinson), respectively. The cells were incubated for 4h in the presence of brefeldin A (GolgiPlug, BD Biosciences, 1:1000 concentration), monensin (GolgiStop, BD Biosciences, 1:1500 concentration) and CD107a antibody. After incubation, normal surface and intracellular staining was performed with the addition of IFNγ and TNF staining. For the flow cytometry based killing assays, NK cells were seeded at a 1:1 effector:target ratio with K562 cells in complete RPMI medium together with Caspase 3-FITC in a 96-well V-bottom plate. The plate was centrifuged at 300 rpm for 3 min and then incubated for 4h at 37°C. Post incubation, the cells were surface stained (CD3-PC5, CD56-ECD, DCM Aqua) followed by fixation.

### Inhibitor experiments

After 4 days of stimulation in 5ng/mL of IL-15, cells were treated either with DMSO, Rapamycin (25 or 50 nM; Sigma) or Torin1 (125 or 250 nM; Apexbio) in the presence of continued IL-15. After 48h of treatment, cells were harvest and stained for flow cytometry analysis.

### Single-cell RNA sequencing

PBMCs from healthy blood donors were screened to determine the educated KIR subset. PBMC were isolated using density gradient centrifugation, followed by NK cell isolation using the DepleteS program on the AutoMACS (Miltenyi Biotec). Isolated NK cells were labelled using 1μM CTV and rested overnight in complete RPMI medium (10% Fetal calf serum, 2mM L-glutamine). CTV-labelled NK cells were stained with the sorting panel and two populations (CD56^dim^NKG2A^+^non-self/selfKIR^-^CD57^-^ & CD56^dim^NKG2A^-^non-selfKIR^-^selfKIR^+^CD57^+^) were sorted for single-cell RNA sequencing while a third population (CD56^dim^ NKG2A^-^ non-selfKIR^-^ selfKIR^+^ CD57^-^) was sorted for culture. Cell sorting was performed at 4°C using a FACSAriaII. Sorted cells were cultured in SCGM (10% human serum, 2mM L-glutamine, 5ng/mL IL-15) in 96 U-bottom wells (200 μL) at 37°C and 5% CO_2_. IL-15 was replenished daily and 100μL of medium was replaced with fresh medium on day 3 of culture. After 6 days of culture, NK cells were stained and FACS sorted based on cell division (CTV) into slowly (CD56^+^ Generation 0-1) and rapidly cycling (CD56^+^ Generation 2+) cells for single-cell RNA sequencing. For single-cell RNA sequencing, 12,000 cells were sorted for each sample (two at baseline, one at day 6) into Eppendorf tubes at 4°C. Cells were washed in PBS + 0.05% BSA and counted. 10,000 cells were resuspended in 35 μL PBS + 0.05% BSA and immediately further processed at the Genomics Core Facility at Oslo University Hospital using the Chromium Single Cell 3’ Library & Gel Bead Kit v2 and the Chromium Controller System (10X Genomics). The sequencing libraries were generated following the recommended protocol. Sequencing was performed on a NextSeq500 (Illumina) with 5∼ % PhiX as spike-inn. Sequencing raw data were converted into fastq files by running the Illumina’s bcl2fastq v2.

### Bioinformatics analysis of single-cell data

The processed 10x files were analyzed in R using the Seurat package. The samples for each time point were merged into one Seurat object (2 samples (1 donor) at baseline, 4 samples (2 donors) at day 6) with retention of the original sample and donor ID. Quality control was performed to remove potential multiplets and to remove outliers based on nUMI and nGene expression. In total 5652 cells passed the quality control phase for the baseline samples and 7556 cells passed the quality control phase for the day 6 samples.

The data was then normalized using a scale factor of 10,000. Next, variable genes were detected using a low x and y cutoff of 0.125 and 0.5 respectively. The dataset was then scaled and regressed on nUMI and nGene. Linear dimensionality reduction was performed using Principle Component Analysis (PCA) on variable genes and the statistically significant principle components (PCs) were determined using both JackStraw and Elbow plots. Using the statistically significant PCs, the cells were clustered using a resolution of 0.9 and visualized using tSNE. Cell cycle scoring was performed using the expression of 94 cell cycle genes. Differentially expressed genes (log(fold change) > 1.2) were then calculated between individual chosen clusters based on sample ID (subset or cycling speed) and cell cycle scoring (cell cycle phase). Heatmaps were used to visualize scaled expression of the differentially expressed genes ordered by log_2_(fold change) in the selected clusters. Clustered heatmaps were used to visualize similarity between cells (annotated by sample ID and cell cycle scoring) in terms of their expression of a given gene set.

Using gene over-expression analysis (Gene Ontology, PANTHER) of the 130 differentially expressed genes at day 6, statistically significantly differences in terms of molecular function, biological processes and pathways were determined. Lastly, using the same 130 differentially expressed genes upstream regulators were determined using Ingenuity Pathway Analysis (Qiagen).

### Statistical analysis

Significance was calculated using either a Wilcoxon signed rank test or when comparing more than 3 groups, a Friedman test followed by Dunn’s multiple comparisons test. Linear regression analysis was performed followed by a Spearman r test to determine significance for correlations. p-values: * <0.05, ** <0.01, *** < 0.001, **** <0.0001. Analysis was performed using GraphPad Prism 6 software.

## Funding

This work was supported by grants from the Swedish Research Council, the Swedish Children’s Cancer Society, the Swedish Cancer Society, the Tobias Foundation, the Karolinska Institutet, the Wenner-Gren Foundation, the Norwegian Cancer Society, the Norwegian Research Council, the South-Eastern Norway Regional Health Authority and the KG Jebsen Center for Cancer Immunotherapy. SP was supported by BB/N01524X/1 from the BBSRC and BJ was funded by a Mildred Scheel postdoctoral scholarship from the Dr. Mildred Scheel Foundation for Cancer Research of the German Cancer Aid Organization.

## Author contributions

A.P. and B.J. conducted experiments. AP analyzed the data. S.L. performed RNA sequencing and generated the libraries. E.H.A. generated the tSNE plots. J.P.G, T.C, E.S. and K.-J.M. provided scientific input. A.P., E.S., and K.-J.M. designed research and wrote the manuscript.

## Competing Interests

KJ Malmberg is a scientific advisor and consultant at Fate Therapeutics.

## Data and materials availability

All data generated and/or analysed during the current study are available from the corresponding author on reasonable request. Due to the current implementation of the Norwegian data legislation at the Oslo University Hospital the RNA Seq datasets of anonymous donors are available upon request in the form of normalized FKPM values.

