## Supplementary Figures for "Intra-lineage Plasticity and Functional Reprogramming Maintain Natural Killer Cell Repertoire Diversity"

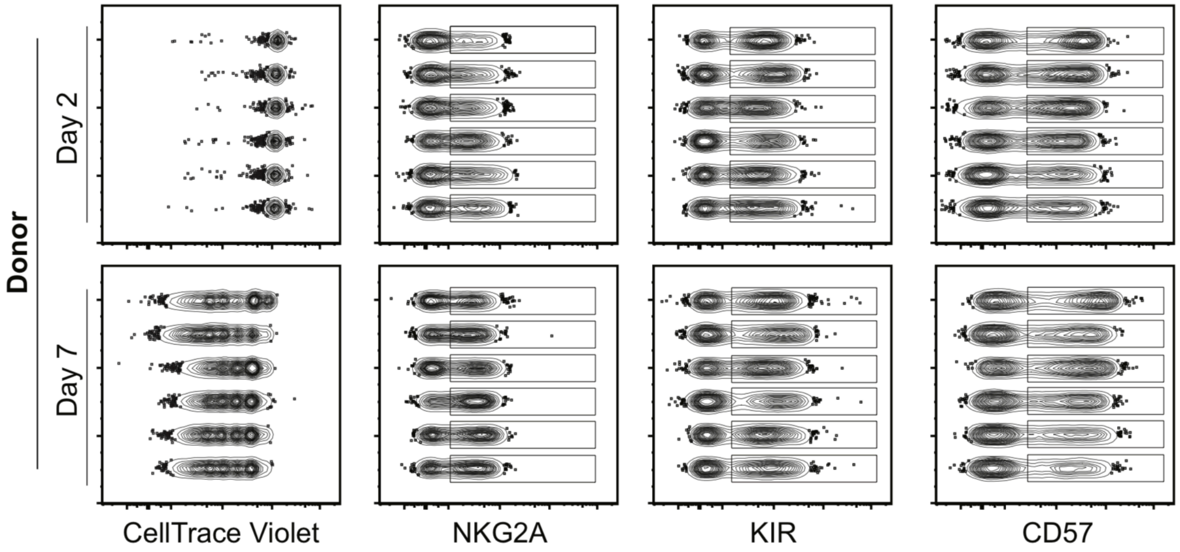


**Fig. S1**. **Inter-donor variation in response to IL-15 stimulation**

Concatenated FACS plots showing CTV dilution and surface expression of NKG2A, KIR and CD57 of the total NK cell population on day 2 and day 7 in IL-15 stimulated cells. n = 6.

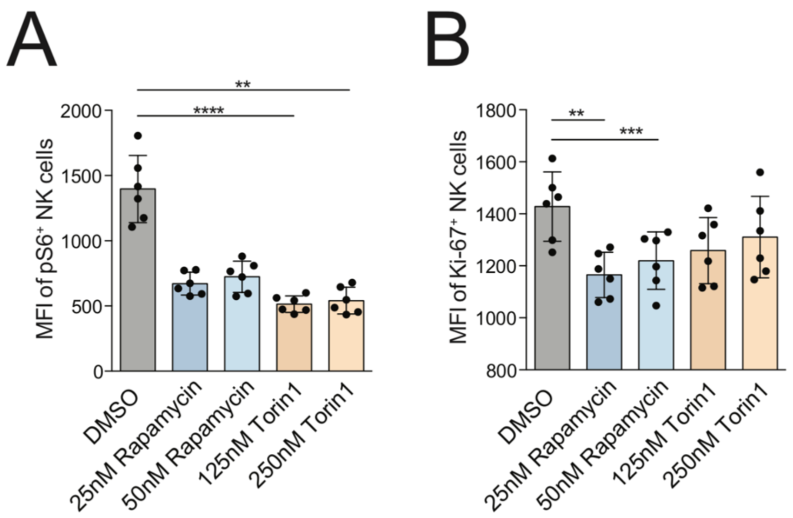


**Fig. S2**. **mTOR inhibition in proliferating NK cells**

(**A** and **B**) Mean fluorescent intensity of pS6 (A) and Ki-67 (B) on day 6 in IL-15 stimulated NK cells treated with DMSO, 25nM Rapamycin, 50nM Rapamycin, 125nM Torin-1 or 250nM Torin-1 for 48h prior to readout. n = 6. Significance was calculated using a Friedman test followed by Dunn's multiple comparisons test (A-B). p-values: * <0.05, ** <0.01, *** < 0.001, **** <0.0001.

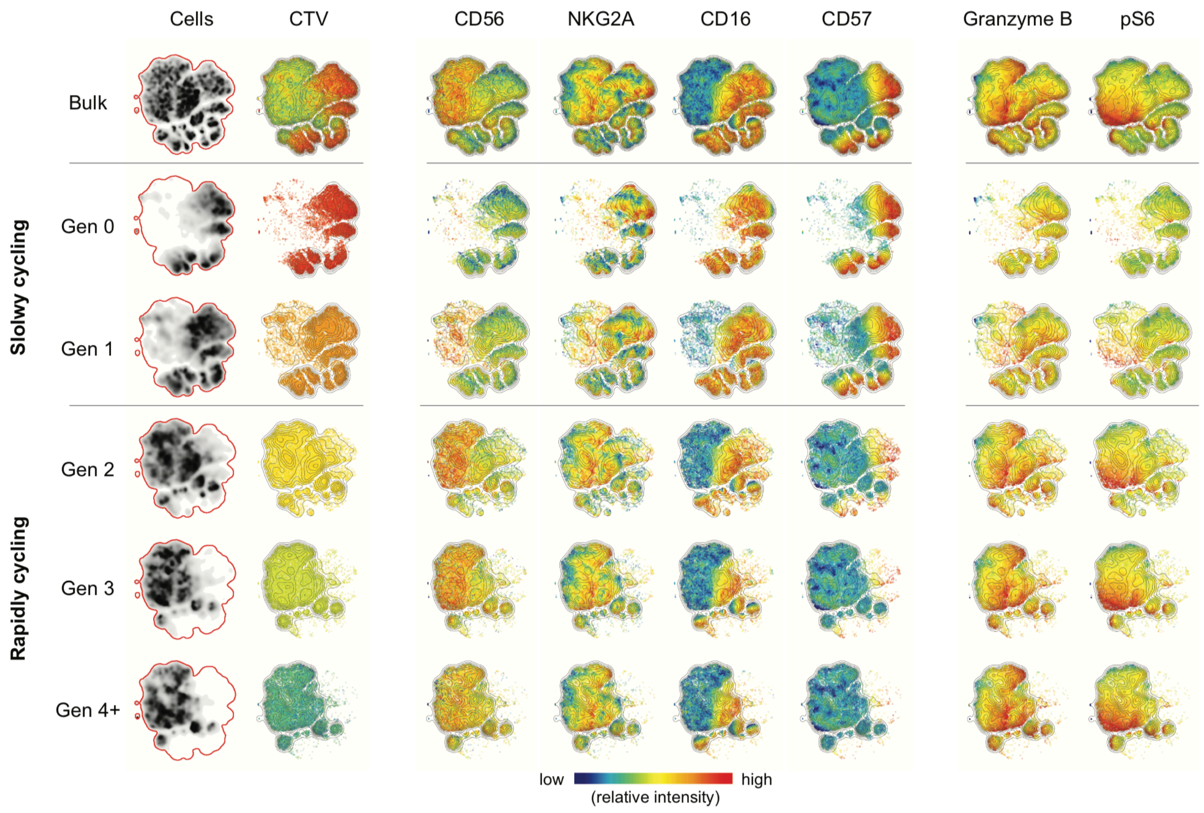


**Fig. S3**. **Phenotype of proliferating NK cells observed at the generation level**

t-SNE plots showing the relative intensity of CD56, NKG2A, CD16, CD57, Granzyme B and pS6 expression in the total NK cell population and gated on individual generations using CTV dilution in one representative donor on day 5. n = 1.

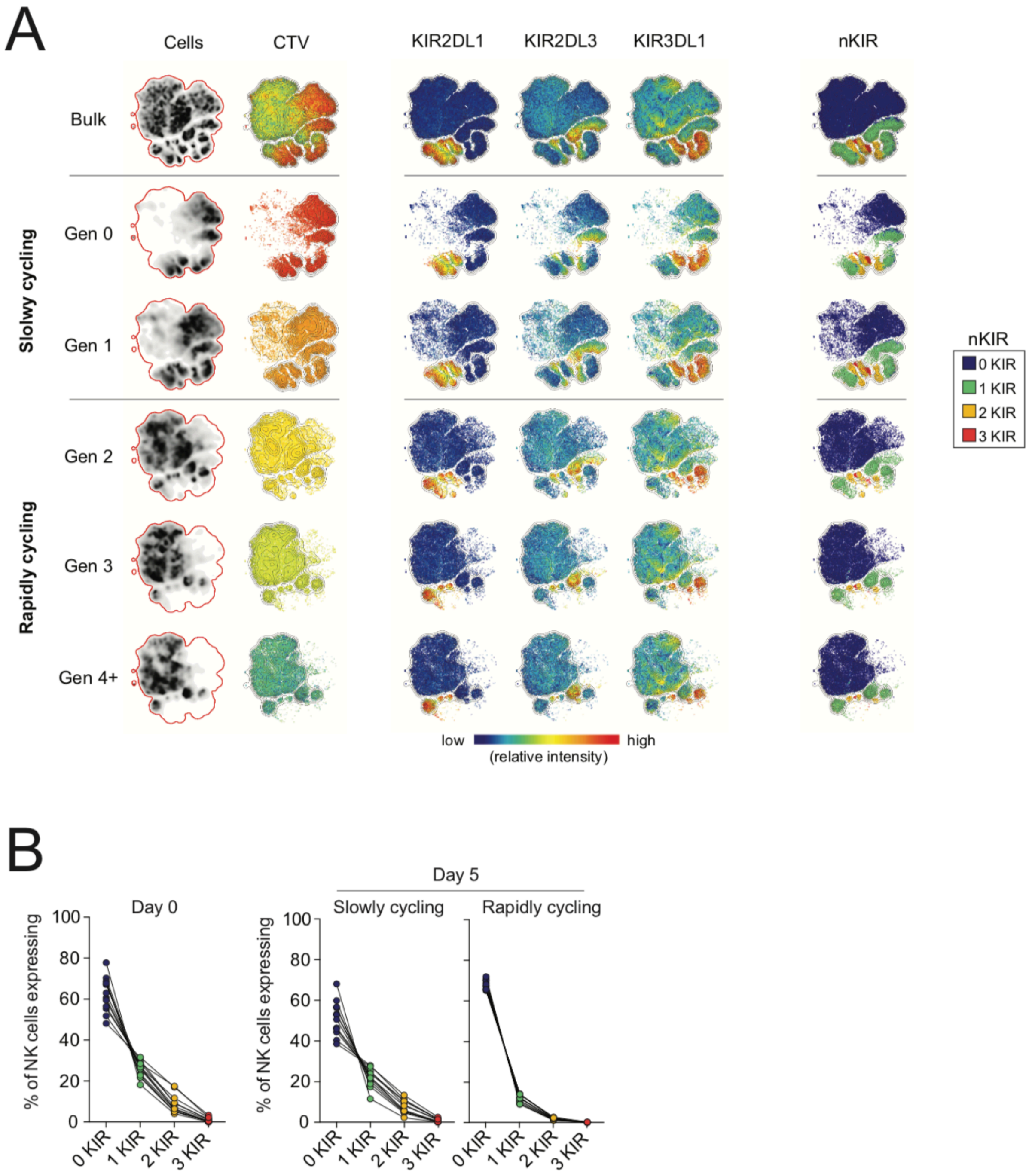


**Fig. S4**. **KIR repertoires of proliferating NK cells observed at the generation level**

(**A**) t-SNE plots showing the relative intensity of KIR2DL1, KIR2DL3 and KIR3DL1 expression, as well as the number of KIR (nKIR) expressed per cell, in the total NK cell population and gated on individual generations using CTV dilutions in one representative donor on day 5. (**B**) The frequency of NK cells expressing 0-3 KIR at baseline and within slowly (Generation 0-1) and rapidly cycling (Generation 2+) cells on day 5. n = 1-12.

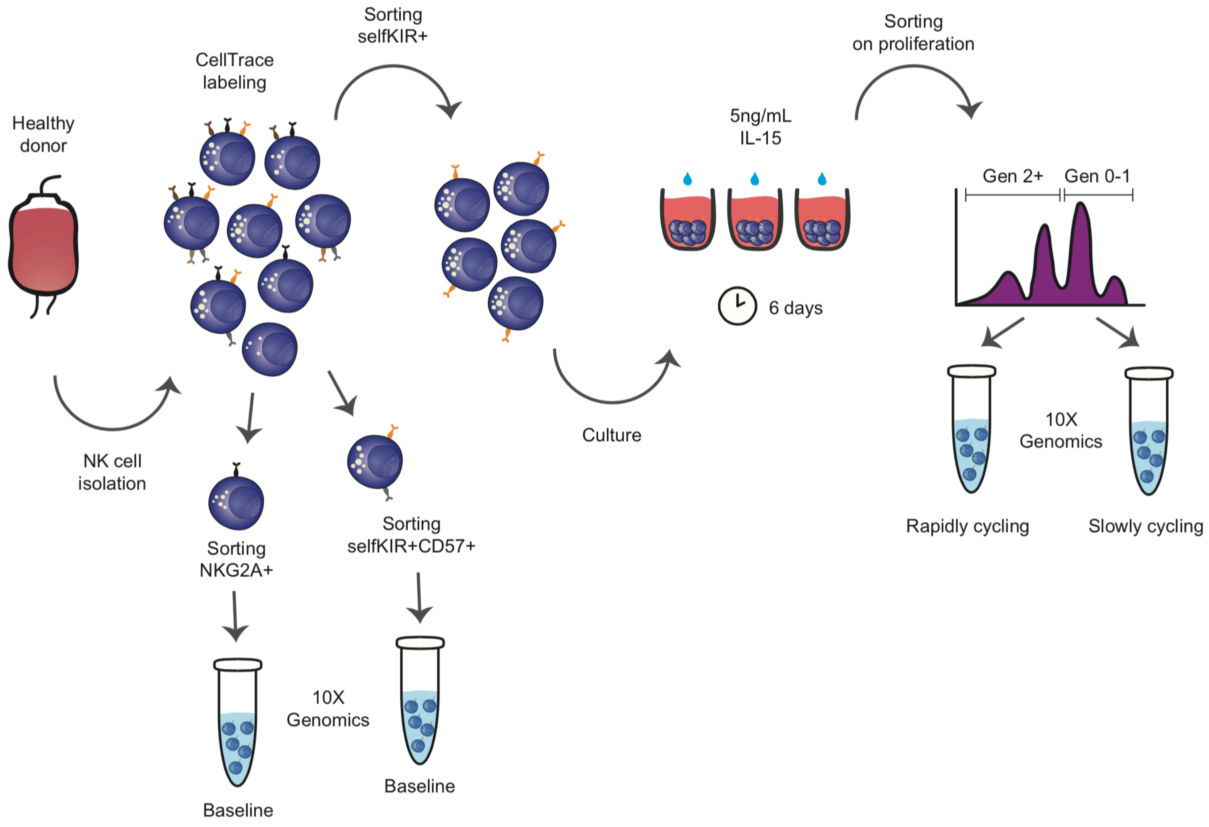


**Fig. S5. Design of single-cell RNA sequencing experiments**

Graphical methodology outline for upstream sample preparation before single-cell RNA sequencing using the 10x Genomics platform.

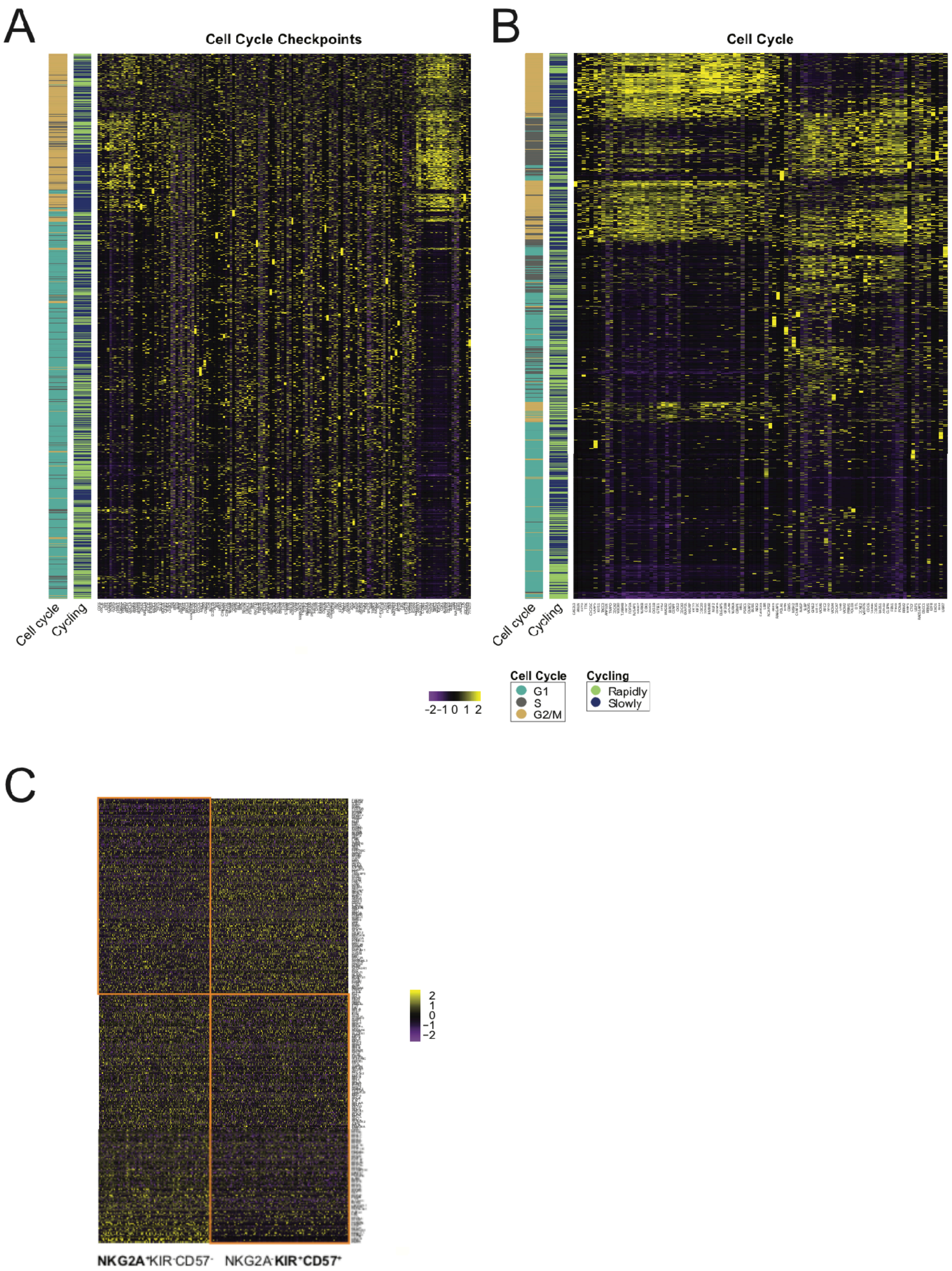
A
Cell Cycle Checkpoints
B
Cell Cycle
Cell cycle
Cycling
Cell cycle
Cycling
-2-1 0 1 2
Cell Cycle
G1
S
G2/M
Cycling
Rapidly
Slowly
C
2
1
0
−1
−2
NKG2A+KIR-CD57-
NKG2A-KIR+CD57+

**Fig. S6**. **Heatmaps of single-cell RNA sequencing data at baseline and in proliferating NK cells.** (**A** and **B**) Expression heatmap for 158 cell cycle checkpoint genes (A) and 94 cell cycle genes (B) shown for all 7556 analyzed cells at day 6. Annotations bars depict the ID of each cell in terms of cycling speed (slowly vs rapidly) and cell cycle phase (G1, S, G2/M). (**C**) Expression heatmap of 295 differentially expressed genes between clusters 1+6 representing NKG2A$^{+}$KIR$^{-}$CD57$^{-}$ cells and clusters 4+7 representing NKG2A$^{-}$KIR$^{+}$CD57$^{+}$ cells from Figure 5D plotted for all 5652 cells analyzed at baseline. n = 1-2.
